# A statistical genetics guide to identifying HLA alleles driving complex disease

**DOI:** 10.1101/2022.08.24.504550

**Authors:** Saori Sakaue, Saisriram Gurajala, Michelle Curtis, Yang Luo, Wanson Choi, Kazuyoshi Ishigaki, Joyce B. Kang, Laurie Rumker, Aaron J. Deutsch, Sebastian Schönherr, Lukas Forer, Jonathon LeFaive, Christian Fuchsberger, Buhm Han, Tobias L. Lenz, Paul I. W. de Bakker, Albert V. Smith, Soumya Raychaudhuri

## Abstract

The human leukocyte antigen (HLA) locus is associated with more human complex diseases than any other locus. In many diseases it explains more heritability than all other known loci combined. Investigators have now demonstrated the accuracy of *in silico* HLA imputation methods. These approaches enable rapid and accurate estimation of HLA alleles in the millions of individuals that are already genotyped on microarrays. HLA imputation has been used to define causal variation in autoimmune diseases, such as type I diabetes, and infectious diseases, such as HIV infection control. However, there are few guidelines on performing HLA imputation, association testing, and fine-mapping. Here, we present comprehensive statistical genetics guide to impute HLA alleles from genotype data. We provide detailed protocols, including standard quality control measures for input genotyping data and describe options to impute HLA alleles and amino acids including a web-based Michigan Imputation Server. We updated the HLA imputation reference panel representing global populations (African, East Asian, European and Latino) available at the Michigan Imputation Server (*n* = 20,349) and achived high imputation accuracy (mean dosage correlation *r* = 0.981). We finally offer best practice recommendations to conduct association tests in order to define the alleles, amino acids, and haplotypes affecting human traits. This protocol will be broadly applicable to the large-scale genotyping data and contribute to defining the role of *HLA* in human diseases across global populations.

## Introduction

More than 50 years ago, some of the earliest complex human disease genetic associations were reported within the major histocompatibility complex (MHC) locus^1,2^. This locus has since been mapped to the short arm of chromosome 6. Sequencing of the human genome has revealed that the MHC locus consists of a cluster of more than 200 genes, including many with immune functions^3^. The MHC locus is broadly divided into three subclasses: the class I region (e.g., *HLA-A, HLA-B and HLA-C* genes), the class II region (e.g., *HLA-DPA1, HLA-DPB1, HLA-DQA1, HLA-DQA2, HLA-DQB1, HLA-DQB2, HLA-DRA, HLA-DRB1, HLA-DRB2, HLA-DRB3, HLA-DRB4* and *HLA-DRB5* genes), and the class III region, which contains additional genes implicated in immune and inflammatory responses (e.g., complement genes)^4^ (**Figure 1a**). Those *HLA* class I and II genes encode protein molecules that form complexes that present antigenic peptides to T cells, thereby influencing thymic selection and T cell activation^4^ (**Figure 1b**). The functional importance of the *HLA* genes and the highly polymorphic nature of this locus have made the MHC region confer the largest number of disease associations of any locus genome-wide (**Figure 1c**). MHC-disease risk is modulated by several underlying mechanisms. For example, in rheumatoid arthritis, polymorphisms in the amino acid sequence of *HLA-DRB1* change the capability of presenting autoantigens^5^ or increase the autoreactive T cells during thymic selection^6^. In another example, the HLA-C*06:02 allele is associated with psoriasis, probably due to increased CD8+ T-cell mediated inflammatory reactions^7^. In another example, schizophrenia’s association within MHC locus was explained in part by structural variation in *C4*, which might modulate synaptic elimination during development^8^.

**Figure 1.**
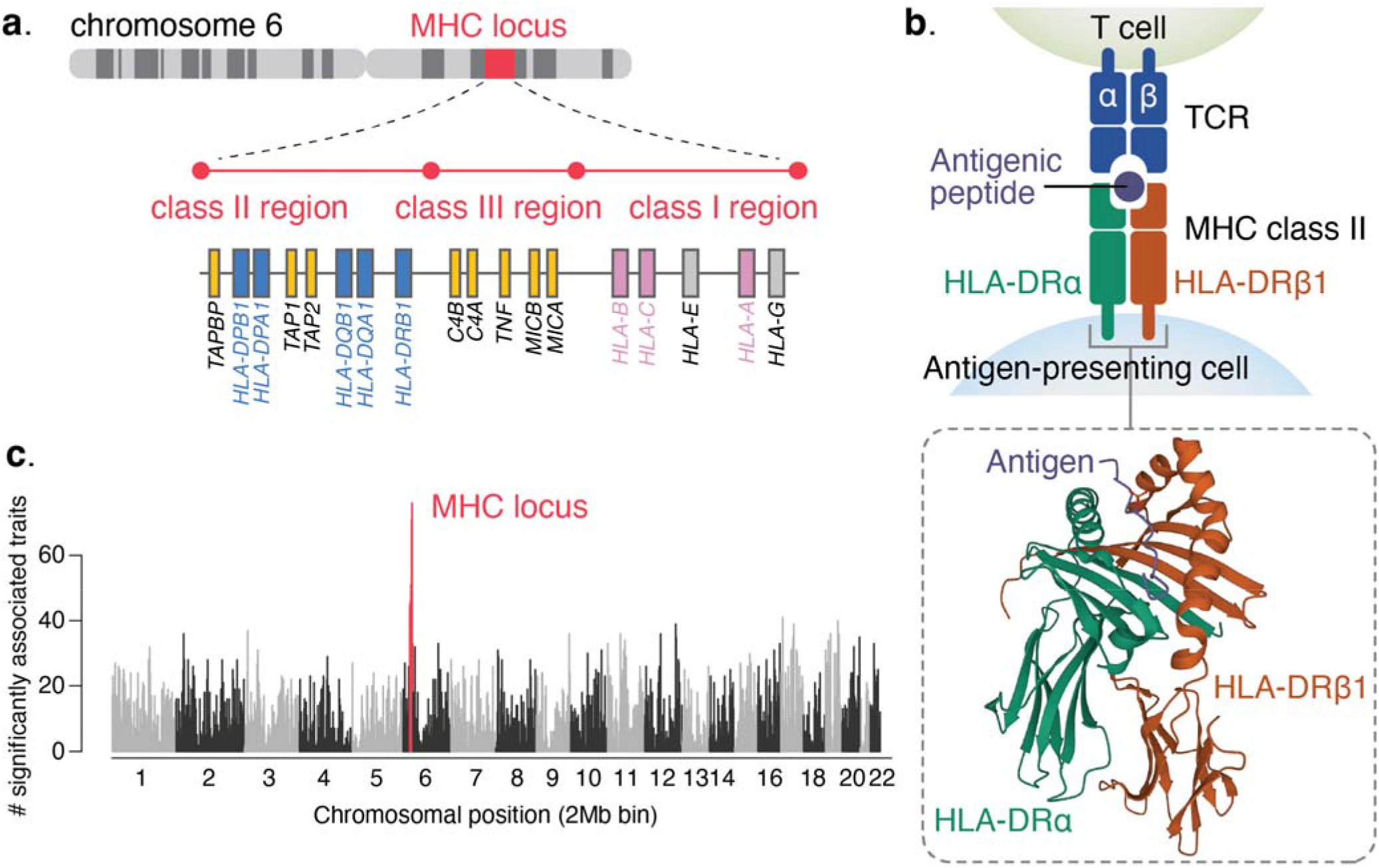
A simplified summary of the location and structure of *HLA* genes on human chromosome 6, and their associations with human traits. **a**. A schematic representation of the human MHC locus, three classes of the region, and genes within them. The genes in pink are the classical class I *HLA* genes, whereas those in blue are the classical class II *HLA* genes. **b**. Presentation of antigenic peptide by an antigen-presenting cell to a T cell through interaction between MHC class II molecule and T cell receptor (TCR). The inset describes protein structure of MHC class II, HLA-DRA and DRB1 adapted from PDB (3L6F). **c**. The number of traits associated with any variants within 2Mb genomic window with *P* < 5×10^−8^ in UK Biobank or meta-analysis of UK Biobank and FinnGen among 198 diseases and biomarkers^9^.

The *HLA* genes within the MHC have been difficult to study because of their highly polymorphic nature, the region’s complex relationship with natural selection, and its unique long-range linkage disequilibrium (LD) structure. The highly polymorphic nature of *HLA* genes renders traditional probe-based genotyping to be challenging. In addition, the genetic diversity at HLA genes is highly population-specific, necessitating efforts to accurately genotype HLA alleles and investigate phenotypic associations in global populations.

These challenges have driven high interest in the genetics community to develop and deploy statistical techniques for HLA alleles. While the direct typing of HLA alleles continues to be costly, labor-intensive and unscalable, *in silico* HLA imputation has recently enabled rapid and accurate estimation of HLA alleles in individuals already genotyped on microarrays. However, there are few guidelines for HLA imputation and to estimate and fine-mapping; these methods are necessary to define HLA effects on human diseases, especially in biobank-scale data from multiple populations.

In this context, here we provide detailed guidelines for imputing HLA alleles and testing for an association with human diseases and traits, in large-scale cohorts and global biobanks. We also provide a step-by-step online tutorial with scripts and available software (https://github.com/immunogenomics/HLA_analyses_tutorial). Definitions of key terms used throughout this article can be found in **Box 1**.

### Box 1

**Key terms and definitions**

|  |  |
| --- | --- |
| MHC region | A genomic region that harbors the major histocompatibility complex (MHC). In GRCh37, it corresponds to chr6:28,477,797-33,448,354 (6p22.1-21.3). |
| Linkage disequilibrium | A non-random association or dependence of alleles at different loci in a given population, making the frequencies of the alleles deviate from the expected when the alleles were independent. |
| Imputation | A procedure of estimating the missing genotypes at loci that are not assayed in the target dataset. |
| Reference panel | A panel of densely genotyped haplotypes to be referred to when predicting the missing genotypes in the target cohort through imputation. |
| Haplotype | A stretch of DNA sequences (including multiple polymorphic loci) along one chromosome that tend to be inherited together due to linkage disequilibrium. |
| Allele | One of two versions of DNA sequences. An individual inherits two alleles (maternal and paternal) for any genomic location. |
| HLA allele | One of the sequence variations at a given *HLA* gene. |
| Genotype | An individual's pattern of DNA sequence at a given location. Two alleles from a mother and a father comprise a genotype. |
| Fine-mapping | A procedure to narrow down and define potentially causal genetic variation(s) affecting the trait of interest, from all the associated genetic variations at a given locus in GWAS by using statistical methods. |
| Homozygous | A state where the two alleles at the genetic variation of interest (e.g., an *HLA* gene) are the same. |
| Heterozygous | A state where the two alleles at the genetic variation of interest (e.g., an *HLA* gene) are different. |
| Allele divergence | A proxy for the functional difference in antigen binding between two *HLA* alleles based on the divergence of their amino acid sequence. |

### Summary of the protocol

The protocol is summarized in **Figure 2a**. The protocol is comprised of two sections: HLA imputation (**Figure 2a-1**) and HLA association testing (**Figure 2a-2**). HLA imputation is a method to infer HLA alleles, amino acids and SNPs from microarray-based genotype within the MHC region. We first introduce the concept of the HLA reference panel (1), which is used as a dictionary to search for similar haplotypes (keyword) to infer unknown HLA types (definition). We highlight specifically our multi-ancestry HLA reference panel, which we recently constructed to enable accurate HLA inference in diverse global populations^10^. We next provide specific instructions to perform QC of the input genotype data (2), per-individual and per-variant (3). The quality of genotype data is critical in achieving accurate imputation, and a special caution should be taken given the extremely complex and polymorphic nature of genetic variants within MHC. We then introduce options to impute HLA (4), either (i) on a user’s local server or (ii) or by using the Michigan Imputation Server (MIS)^11^, which is a publicly available, web-based imputation platform we jointly support with Michigan University. We finally describe the quality metrics and post QC of the imputed variants (5).

**Figure 2.**
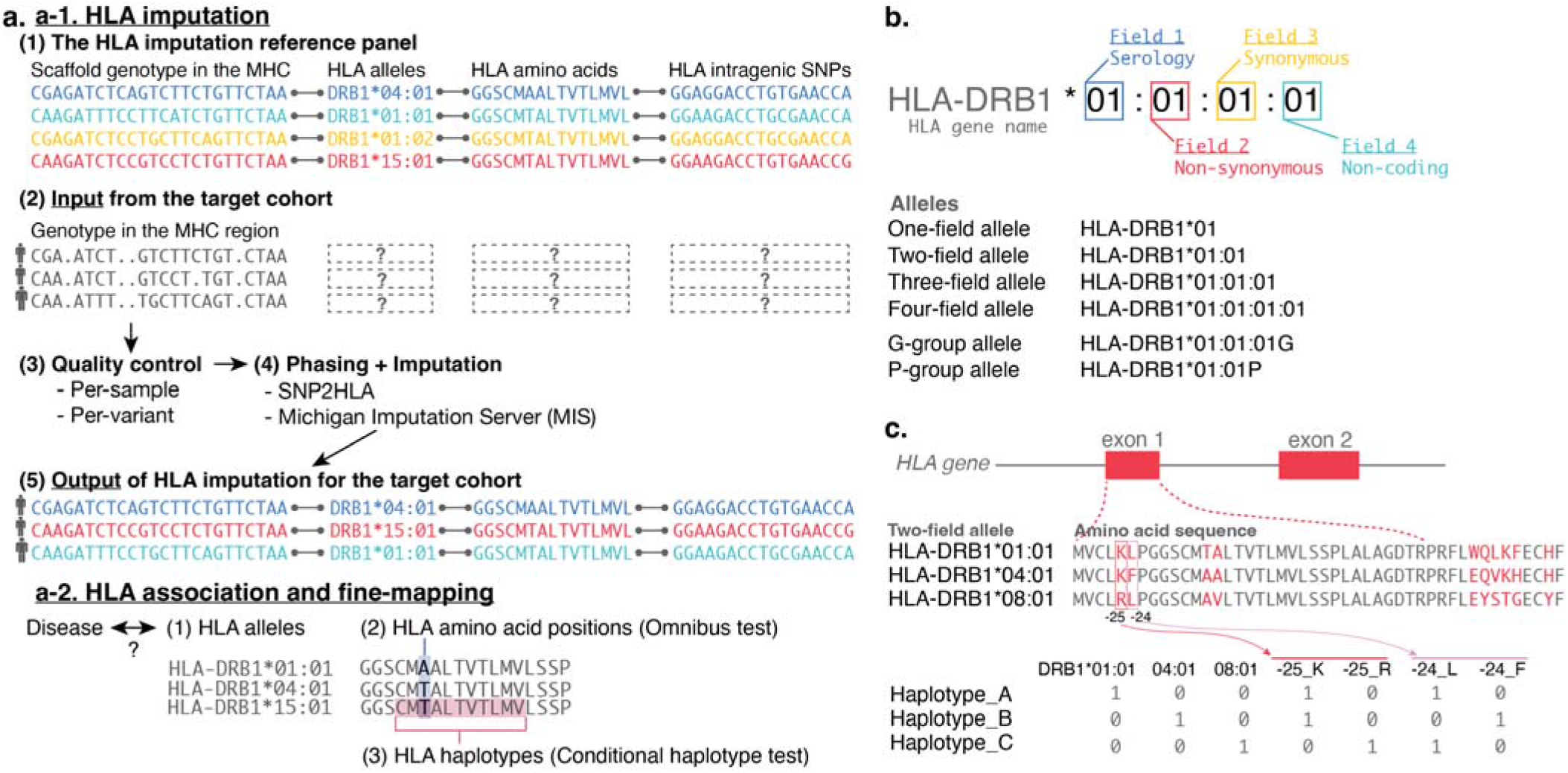
The overview of HLA imputation, association, and fine-mapping, including construction of HLA reference panel. **a**. Overview of this protocol. **(a-1)** A toy example of HLA imputation, describing (1) HLA imputation reference panel, (2) input genotype in the MHC region from the target cohort without HLA types, (3) quality control of the target genotype, (4) genotype phasing and imputation to predict the untyped HLA alleles in the target cohort, and (5) output of the predicted HLA alleles. **(a-2)** Statistical methods to investigate and fine-map association of HLA alleles, amino acids and their haplotypes with a trait of interest. **b**. The naming system (nomenclature) of HLA alleles, consisting of four fields with each field corresponding to the types and consequences of nucleotide variations. **c**. (top) The amino acid sequences defining each of three example HLA-DRB1 alleles. The amino acids colored in red indicate the positions where they have variations among the alleles. The numbers (−25 and -24) at the bottom indicate the relative position of those amino acids within a coding region of HLA-DRB1. (bottom) A procedure to code each of the HLA alleles and amino acid polymorphisms as binary markers: 1 if that marker is present within a haplotype and 0 otherwise. Each of the residues are coded separately for a given amino acid position in the corresponding HLA protein.

We next describe statistical methods to perform comprehensive association tests between HLA genotype and human traits (**Figure 2a-2**). Since HLA associations are often explained by amino acid sequences in the peptide binding groove of HLA molecules^12^, we describe strategies to fine-map associations with the aim of pinpointing causal variation. We start from a simple single-marker test which is similar to that commonly used in GWAS, and then elaborate on the HLA-specific fine-mapping methods (e.g., an omnibus test (2) and a conditional haplotype test (3)). We also introduce statistical tests to define non-additive, interactive, and multi-trait contribution of HLA alleles.

### Introduction to HLA nomenclature

Sequence variation within *HLA* genes is organized by the International Immunogenetics database (IMGT)^13^, which has documented and named 33,490 unique HLA alleles (URL: https://www.ebi.ac.uk/ipd/imgt/hla/about/statistics/). Within each of the HLA alleles, there are nucleotide variants which sometimes cause amino acid changes (i.e., non-synonymous nucleotide substitutions) and sometimes not (i.e., synonymous, intronic and intergenic nucleotide substitutions). A detailed nomenclature system at IMGT has been developed to organize those polymorphisms in *HLA* genes into four fields (**Figure 2b**)^14^. In this nomenclature, field 1 (i.e., the first two digits, e.g., HLA-DRB1*<u>01</u>) describes the serological type, which was historically defined based on similar seroreactivity to immunological reagents. Field 2 (i.e., the next set of digits, e.g., HLA-DRB1*01:<u>01</u>) corresponds to the unique amino acid sequence of the gene; all the non-synonymous changes are reflected in this set. Field 3 (e.g., HLA-DRB1*01:01:<u>01</u>) reflects synonymous nucleotide substitutions within the coding sequences, and field 4 (e.g., HLA-DRB1*01:01:01:<u>01</u>) reflects polymorphisms within the intronic or non-coding regions. Thus, whereas nucleotide variants define HLA alleles at up to four-field resolution, most disease associations are captured by two-field HLA resolution since amino acid sequence captures most of the structural differences between the alleles.

The four-field naming system is the current standard and most widely used, but it is worth expanding upon the alternative nomenclatures since they are sometimes seen in practice. Before the current four-field naming system was introduced, the IMGT had used the nomenclature without a field separator (‘:’), where each field must have two digits. Therefore, one-field alleles had been called two-digit alleles, and two-field alleles had been called four-digit alleles. However, as the number of two-field alleles belonging to a given one-field allele began to exceed 100 (e.g., HLA-A*02101 and HLA-B*15101), the name “four-digit” designation became inappropriate. Thus, the IMGT updated the previous nomenclature system by introducing the field separator (e.g., HLA-A*02:101 and HLA-B*15:101) and four-field naming system.

In this same update, the IMGT introduced two additional nomenclature schemes to facilitate practical reporting of HLA typing: G group and P group. Current classical HLA typing technologies sometimes cannot resolve an HLA allele at four-field resolution and define a group of similar alleles based on the variations within peptide binding domains (exon 2 and 3 for class I *HLA* genes and exon 2 for class II *HLA* genes). The G group nomenclature represents HLA alleles that share the same nucleotide sequence in the peptide binding domains. For instance, HLA-A*01:02:01G includes HLA-A*01:02:01:01, HLA-A*01:02:01:02, HLA-A*01:02:01:03, and HLA-A*01:412, but not HLA-A*01:02:02. The P group nomenclature represents HLA alleles that share the same protein sequence in the peptide binding domains. For example, HLA-A*01:02P includes HLA-A*01:02:01:01, HLA-A*01:02:01:02, HLA-A*01:02:01:03, HLA-A*01:02:02, and HLA-A*01:412.

*Introduction to HLA imputation*

Genotype imputation is the term used to describe estimation of missing genotypes that are not assayed in the target dataset. Most imputation methods use data from densely genotyped samples as a reference dataset in which haplotypes have been inferred^15^. They typically use statistical approaches such as hidden Markov models (HMM) to fill in missing genotypes in a dataset of interest with incomplete genotype data. Here the genotype data reflects the observed states, while the template haplotypes are represented as the unknown hidden states. Most imputation algorithms produce a probabilistic prediction of each imputed genotype. These probabilities can be used to either (1) calculate a probabilistic dosage, which is a simple sum of those expected probabilistic allele count, or (2) a best-guess genotype, which is a combination of alleles which have the largest probability. These values can then be used in the downstream analyses. Dosages inferred from imputed results are a continuous value between 0 and 2, whereas guess genotypes are discrete values of 0, 1, or 2 alleles. Genotype imputation can boost the power of the association studies, fine-map the signal, and enable meta-analysis of multiple cohorts^15^.

After imputation, it is essential to understand the accuracy of imputation. The quality of predictions can be technically measured by masking the genotype, imputing them, and deriving the correlation between the true (masked) genotype and the predicted genotype. We favor using this correlation as a metric, as opposed to accuracy (percent of concordance between true genotype and imputed genotype calls), since accuracy can be upwardly biased for rare alleles. In practice, true genotype data is often missing. In these instances, we can also estimate the quality of imputation by the ratio of the empirically observed variance of the allele dosage to the expected binomial variance at Hardy-Weinberg equilibrium (*Rsq*).

HLA imputation is natural extension of the genotype imputation. The HLA imputation infers HLA alleles, amino acid polymorphisms, and intragenic SNPs within *HLA* (hidden state). Due to the excessive variation of these *HLA* genes, these variants generally cannot be accurately assayed with popular probe-based genotyping arrays. HLA alleles are inferred indirectly by using surrounding genotyped SNP variants in the MHC region (“scaffold” variants; **Figure 2c**). Reference haplotypes are constructed from samples with both genotyped SNP variants and HLA alleles genotyped by either classical sequence-based typing (SBT)^16^ or inference from untargeted sequencing data, such as whole genome sequencing (WGS) data^17,18^. The HLA amino acid sequences and intragenic SNPs within *HLA* genes can also be included in the reference haplotypes to enable their imputation. There are many widely used statistical software tools to perform the HLA imputation, such as SNP2HLA^19^, HIBAG^20^, and HLA*IMP^21^, HLA-IMPUTER^22^, and GRIMM^23^. The SNP2HLA and HLA*IMP methods use the same HMM algorithm used in genome-wide imputation, whereas the HIBAG method uses a machine-learning technique: a bagging method^24^. Imputation performance is often related to the size, quality, and suitability of the reference panel rather than the statistical software used. The output of the HLA imputation is a posterior probability as well as an effective dosage (ranging from 0 to 2) for each HLA allele in a given sample. Subsequent association tests usually account for the uncertainty of the imputation by using the estimated dosage as an explanatory variable.

### HLA imputation reference panel

There have been many efforts to construct haplotype reference panels in the MHC region to enable HLA imputation. Since the haplotype structure within the MHC region differs significantly among populations^10^, it is important that the target dataset is well represented by the reference haplotype panel. The current availability of published HLA reference panels is summarized in **Table 1**.

**Table 1.**
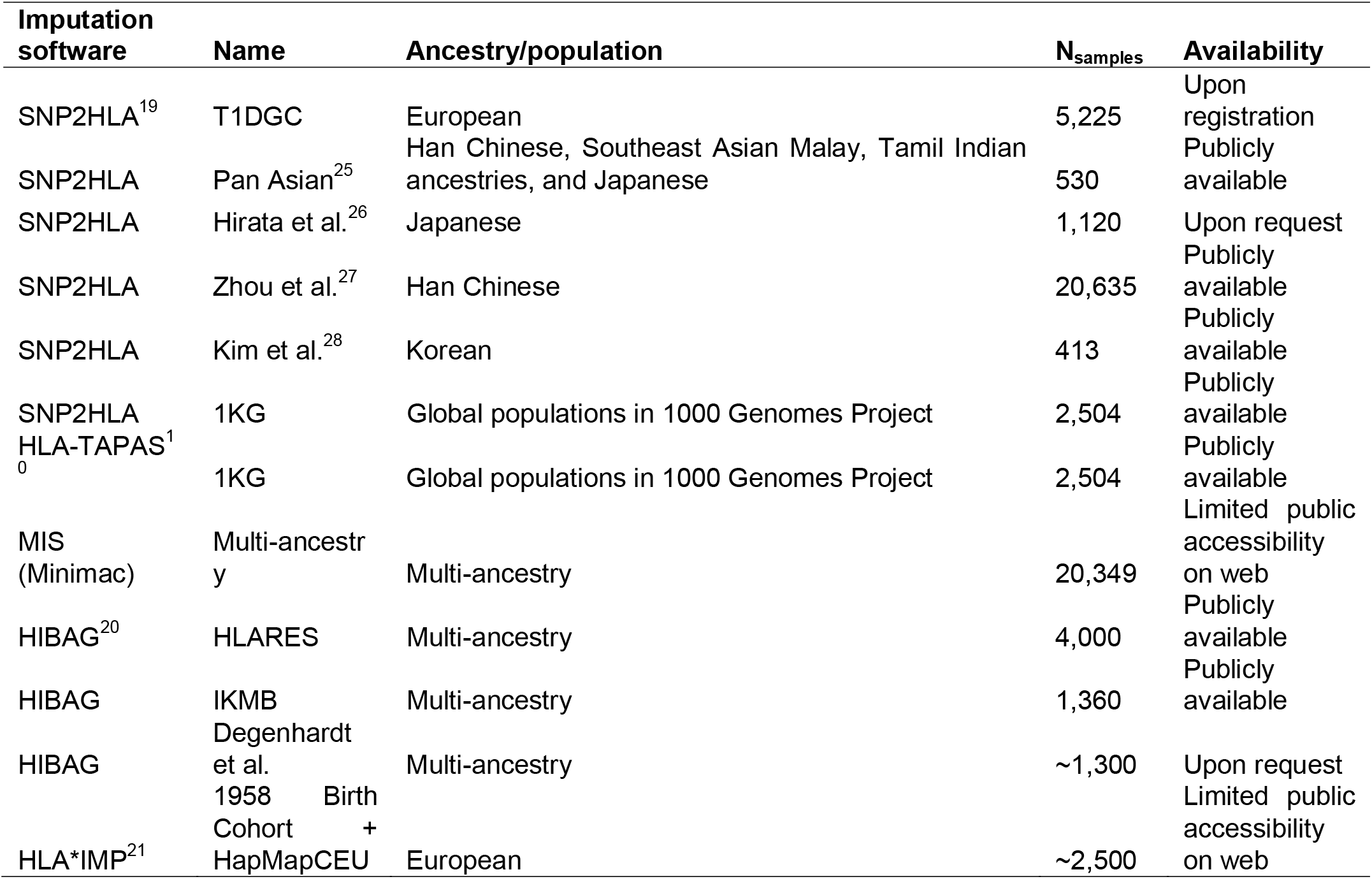
A list of available HLA imputation reference panels.

A list of currently available HLA imputation reference panels, the sample ancestry, the number of samples, and whether they are publicly available or not. Limited public accessibility means that while the raw reference panel (individual-level genetic data) is not accessible, users can use it for imputation via web-based imputation service. MIS: Michigan Imputation Server.

It is also possible to construct a custom HLA reference panel. SNP2HLA and HLA-TAPAS^10,19^ are tools to construct such custom reference panels. Starting with a SNP genotyped cohort (“scaffold variants”), we can either (1) obtain the gold standard SBT of HLA alleles (such as sequence-specific oligonucleotide probe hybridization (SSOP)^16^) if DNA is available or (2) infer HLA alleles from WGS (e.g., HLA*PRG and HLA*LA)^17,18,29^. Reference panels can include alleles of classical *HLA* genes (*n*_gene_ = 8)^10^, which are most polymorphic and disease-associated, or both classical and non-classical *HLA* genes (*n*_gene_ = 33)^26^. In the SNP2HLA algorithm, HLA alleles are converted to biallelic markers (e.g., 1 indicates the presence of the allele and 0 indicates the absence of the allele). Classical SBT, such as SSOP, is the most accurate approach to HLA genotyping. Incorporation of SBT genotypes into reference panels results in highly accurate imputation; however, since SBT is costly and labor-intensive, it cannot be easily used to build large reference panels. Graph-based inference of HLA alleles from WGS is a potential alternative method that can be easily applied to large sequencing datasets that are increasingly available^17,18,29^. However, an important caveat is that the accuracy of HLA typing by those graph-based methods can be variable. For example, imputation performance is affected by (i) quality of the sequencing data, (ii) read depth and length, (iii) representation of the population in reference databases such as IMGT, and (iv) the degree of sequence variation within the targeted *HLA* gene. For studying under-represented populations or highly polymorphic genes, gold standard SSOP might still be necessary to construct a suitably accurate reference panel.

To enable imputation of amino acid polymorphisms and intragenic HLA SNPs, we can encode all these variants as binary markers based on the refence amino acid and nucleotide sequences of each observed HLA allele from the IMGT HLA Database (https://www.ebi.ac.uk/ipd/imgt/hla/) (**Figure 2d**). The scaffold genetic variants within the MHC region are usually obtained by either genotyping with a SNP microarray or WGS. Stringent SNP QC is essential for accurate phasing, and ultimately accurate imputation. In constructing and updating a multi-ancestry HLA reference panel, we optimized this QC process to maximize imputation accuracy. Specifically, we started with QCing each of the global cohorts separately, with genotype call rate (> 95%) and sample call rate (>90%). We then retained all the variants that were present in the 1000 Genomes Project and excluded any variants that were not included in commonly used genotyping arrays (Illumina Multi-Ethnic Genotyping Array, Global Screening Array, OmniExpressExome, and Human Core Exome), since these variants that are not included in the target genotype data are more likely to result in phasing switch errors without improving imputation accuracy. When combining all the cohorts to construct the multi-ancestry panel, we cross-imputed all the variants together to avoid excluding population-specific variants that are polymorphic in a specific cohort but monomorphic and thus not called in the other cohorts (**Supplementary Figure 1**). The final reference panel includes the HLA alleles, amino acids, intragenic HLA SNPs, and the “scaffold” variants (i.e., SNP variants outside of *HLA* gene but within the extended MHC region), which are then phased statistically or by using trios.

Imputed HLA alleles and variants are often used for subsequent association testing and meta-analyses to fine-map disease risk. Such studies potentially include data from multiple cohorts, datasets, or populations. To avoid spurious associations due to batch effects and population stratification, it is essential to perform HLA imputation on all datasets using the same reference panel, ideally with all case and control samples genotyped together. Given that such case-control cohorts may originate from multiple populations to increase the fine-mapping resolution, we previously constructed an HLA reference panel covering multiple global populations^10^.

With the publication of this protocol, we present an updated version of this multi-ancestry panel (version 2). Briefly, we added samples from European (*n* = 2,233) and Japanese (*n* = 723) ancestry for a total of 20,349 individuals. This panel represents admixed African, East Asian, European and Latino populations. We also updated HLA allele calls and a set of scaffold variants. We plan to maintain and update the panel further to increase representation of globally diverse populations, improve the HLA allele calls, and refine selection of the scaffold variants to achieve the most accurate imputation.

### Recommendations for collecting genotype and phenotype information

When designing a study to investigate the effect of HLA variation on human traits, it is important to be strategic when collecting genotype and phenotype data. For genotype data collection, one should ensure that the genotyping array used for the target cohort has a high coverage in the MHC region in order to adequately tag, through LD, HLA alleles, which contributes to accurate imputation. While most currently used genotyping arrays include a sufficient number of SNPs to tag HLA alleles for accurate imputation, some arrays have limited SNP coverage of the MHC region (**Supplementary Table 1**)^30^. We and others have shown that lower MHC coverage results in inaccurate imputation^19,31^. Furthermore, all study participants should ideally be genotyped together with the same genotyping array, to avoid introducing any structure that could cause a bias in imputation and the subsequent association testing and possibly fine-mapping.

Careful phenotype curation is very important when fine-mapping disease-associated variants. Discovery of HLA association signals can be enhanced by the addition of more samples, even at the risk of misclassified samples. However, fine-mapping can be affected by including misclassified samples. For example, studies of autoimmune disease including individuals with different subgroups of patients can obscure efforts to localize disease alleles. This has for instance been observed in rheumatoid arthritis, where patients with positive antibody status are phenotypically and genetically different from those with negative antibody status^32,33^. Recently, many efforts have been made to curate the phenotypes in large-scale biobanks^34^ using self-reported disease status or billing code (e.g., ICD-10)^35^. While the total number of samples with these forms of phenotyping is large in these biobanks and may enable discovery, imprecise phenotype labeling may confuse HLA fine-mapping. In contrast, physician-curated cohorts may be important for fine-mapping efforts.

In addition to disease phenotypes, one must exercise caution when measuring HLA-related molecular phenotypes, such as *HLA* gene and protein expression. It is well established that HLA gene and protein expression is affected by the *cis-*regulatory genetic variants (i.e., expression quantitative trait loci (eQTL) and protein expression quantitative trait loci (pQTL))^36–38^. When conducting eQTL studies, measuring *HLA* expression in RNA-seq is particularly challenging due to the high degree of genetic polymorphism among individuals. Standard expression quantification pipelines rely on a single human reference genome to align sequencing reads. The number of reads mapping to each *HLA* gene might be biased for two reasons: (1) the reads may fail to map to the reference due to the high degree of sequence variation (i.e., a large number of mismatches) and (2) the reads may not uniquely map to a single gene in the reference due to the similarity among nearby *HLA* genes (i.e., multi-mapping)^38^. To address this, more accurate gene expression estimates can be obtained by using an HLA-personalized reference^38^; instead of using a standard single human reference genome, we can supply customized HLA sequences for each target individual for each *HLA* gene (either based on classical HLA typing or HLA imputation) to minimize the degree of variation between the RNA-seq reads and the reference and hence reduce the possibility of mapping failures and multi-mapping. Similarly, caution should be taken for HLA pQTL studies. HLA protein expression is often measured by antibody-based methods (e.g., antibody-derived tags) at single-cell resolution. However, these antibodies may have differing binding affinities to the protein products of different HLA alleles. We should take caution when conducting pQTL studies, since this differing affinity might cause a bias towards specific HLA alleles when measuring the abundance of HLA proteins across individuals.

### Quality control of the target genotype data

Data quality control (QC) of genotype data prior to HLA imputation is extremely important. We next outline the basic QC measures commonly used in GWAS^39^, as well as specific instructions to handle genetic variants within the MHC region. These QC measures are typically performed once for each genotyping batch, followed by data integration and the final QC for the combined dataset (**Figure 3**).

**Figure 3.**
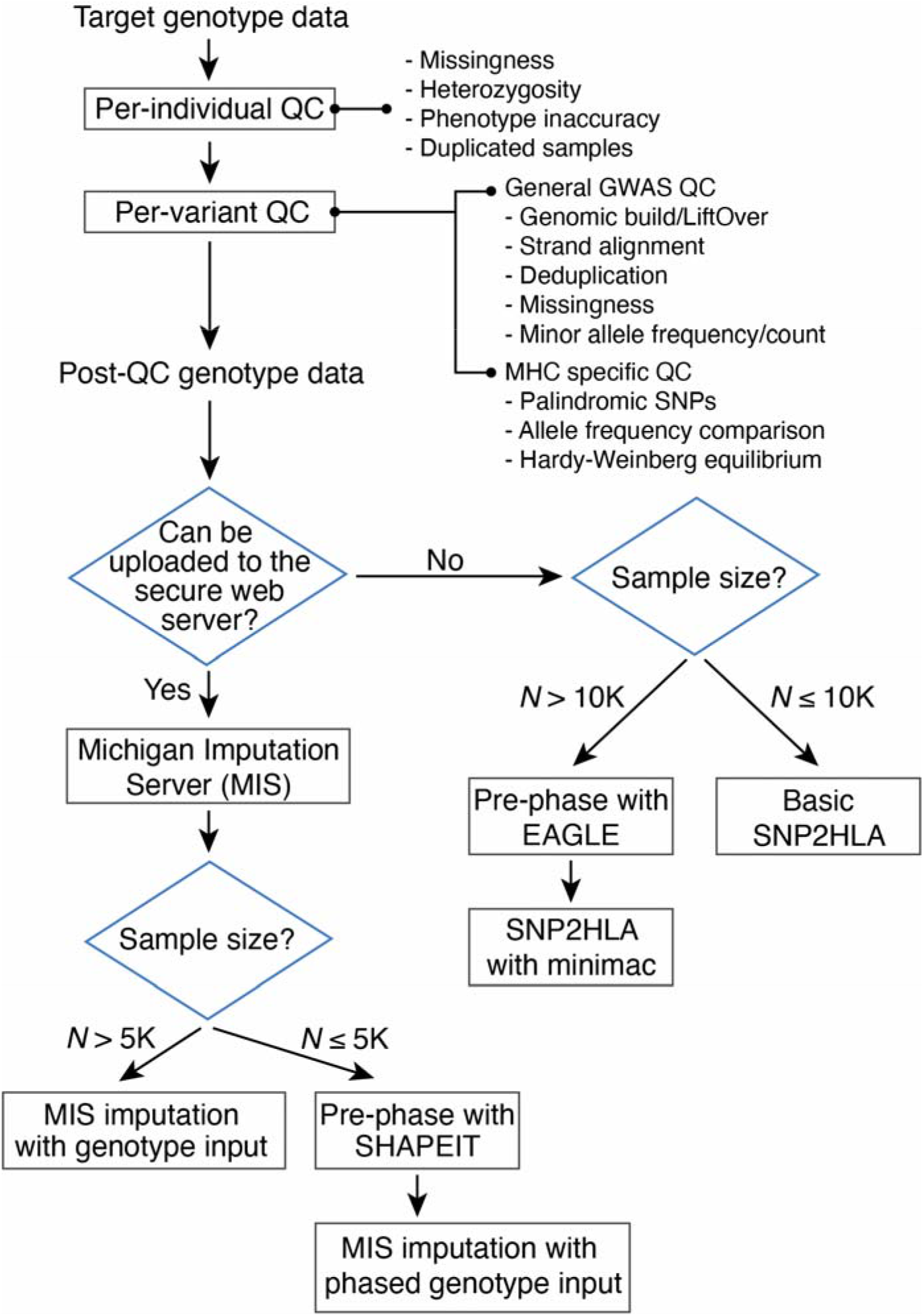
A flow chart of suggested analytical steps for genotype QC and HLA imputation. A best-practice guideline to impute HLA alleles by using SNP2HLA algorithm, depending on the characteristics of the target genotype data.

#### Per-individual QC

We follow established guidelines^34,39,40^ to perform standard per-individual QC in GWAS. Typically, we remove (i) individuals with high missingness (e.g., > 0.02), (ii) individuals with outlier high heterozygosity on suspicion of sample contamination, (iii) individuals with discordant sex information between the meta data and genotype, and (iv) individuals suspected to be duplicate samples based on genotype relatedness. We note that the threshold for each QC measure could be data-dependent, and thus we recommend reviewing the distributions of those metrics for each of the datasets.

#### Per-variant QC

It is important to select high-quality variants to achieve accurate imputation. We will describe the variant QC that is generally recommended for GWAS as well as specific considerations for the MHC region. As part of standard GWAS QC, we recommend ensuring that the target genotype data has genomic positions based on the same genome build as the reference panel. LiftOver software^41^ can be used to lift the genomic position over to the desired genome build. Next, genomic variants are typically aligned to the forward strand to be consistent with the reference panel. We also identify duplicated variants within the dataset based on genomic position and alleles, and de-duplicate them by removing ones with higher missingness. We then remove (i) variants with high missingness (e.g., > 0.01), (ii) variants demonstrating a significant deviation from the Hardy-Weinberg equilibrium (HWE), and (iii) variants with very low minor allele frequency (MAF). Specifically, we remove variants with very low MAF (e.g., < 0.01 or 0.005) or small minor allele count (MAC; e.g., < 5), assuming low accuracy in genotype calling from clustering. The sample size and the estimated ancestry should be accounted for when selecting the threshold in order to retain informative population-specific markers. We usually only keep biallelic variants and remove multi-allelic variants for simplicity in the imputation.

Specific caution should also be taken for per-variant QC in the MHC region, due to (i) highly variable allele frequency (AF) of variants within MHC across populations and (ii) expected HWE deviation in the MHC variants due to natural selection. For example, we usually align target genotype alleles to the forward strand. For non-palindromic SNPs (i.e., SNPs without A/T or G/C allele combinations), it is easy to do so by looking up the alleles with the same position in the reference human genome sequence on forward strand. If the alleles between the target and the reference genome are different (e.g., A/C in the reference but T/G in the target), we flip the alleles in the target dataset (swap alleles from T to A and from G to C in the target). On the other hand, in handling palindromic SNPs (i.e., SNPs with A/T or G/C alleles), we usually compare population-derived AF and the AF in the target dataset to eliminate allele ambiguity. If the AFs between them are largely different (e.g., A: 20% and T: 80% in the population reference but A: 78% and T: 22% in the target), we can flip the alleles to be consistent with the population-derived AF (swap alleles from A to T and from T to A in the target). However, this strategy might be ineffective within the MHC since reference AF for those SNPs might be different from the target samples when the study population is different, when there are large AF differences between cases and controls in case-control studies, or when the study sample size is too small to estimate AF accurately. Therefore, when the strand information of those palindromic SNPs is ambiguous in the target genotyping array or the genotyped data, it may be preferable to exclude all the palindromic SNPs. Second, we may compare AF of the variants after QC in the target data with AF in the population-frequency database (e.g., 1000 Genomes Project^42^ and gnomAD^43^) or AF in the reference panel as a sanity check. When the AFs are very different between the two, those variants could be subject to genotyping error and should probably be removed. However, when the population does not exactly match between the target and the database or the reference, this strategy might be ineffective within the MHC. Thus, we could consider using a liberal threshold when removing variants based on the AF differences. Third, the extreme deviation from HWE is usually indicative of a genotyping or genotype-calling error that results in poor clustering^44–46^ and is used as a metric to exclude poor quality variants. However, the deviation from HWE is to some extent expected in the MHC region due to natural selection^47^ or due to the difference in allele frequency between cases and controls. The expected deviation will be greater when we study a cohort from multiple populations or of admixed ancestry, or when the effect size of HLA on the disease is large. Therefore, for the purpose of per-variant QC, we could consider (1) calculating HWE *P* values only within control individuals (as is generally recommended in GWAS), (2) calculating HWE *P* values within individuals from a representative single-ancestry, or (3) using liberal threshold such as HWE *P* < 1×10^−10^ when removing the variants suspected of poor clustering while retaining the important markers for HLA imputation. When we are unsure about the threshold, an appropriate value can be identified by visually inspecting the genotype cluster plots.

### Tools for genotype phasing and HLA imputation

Once we QC the target genotype data and prepare the optimal HLA reference panel, we start HLA imputation for the target data using existing tools. **Table 2** summarizes the main software programs for HLA imputation and the available HLA reference panels. Of note, some imputation programs take as input the genotype files directly after the QC as described above, while others require users to pre-phase the genotypes to obtain haplotypes^11,21^ before imputation (**Figure 3**).

**Table 2.**
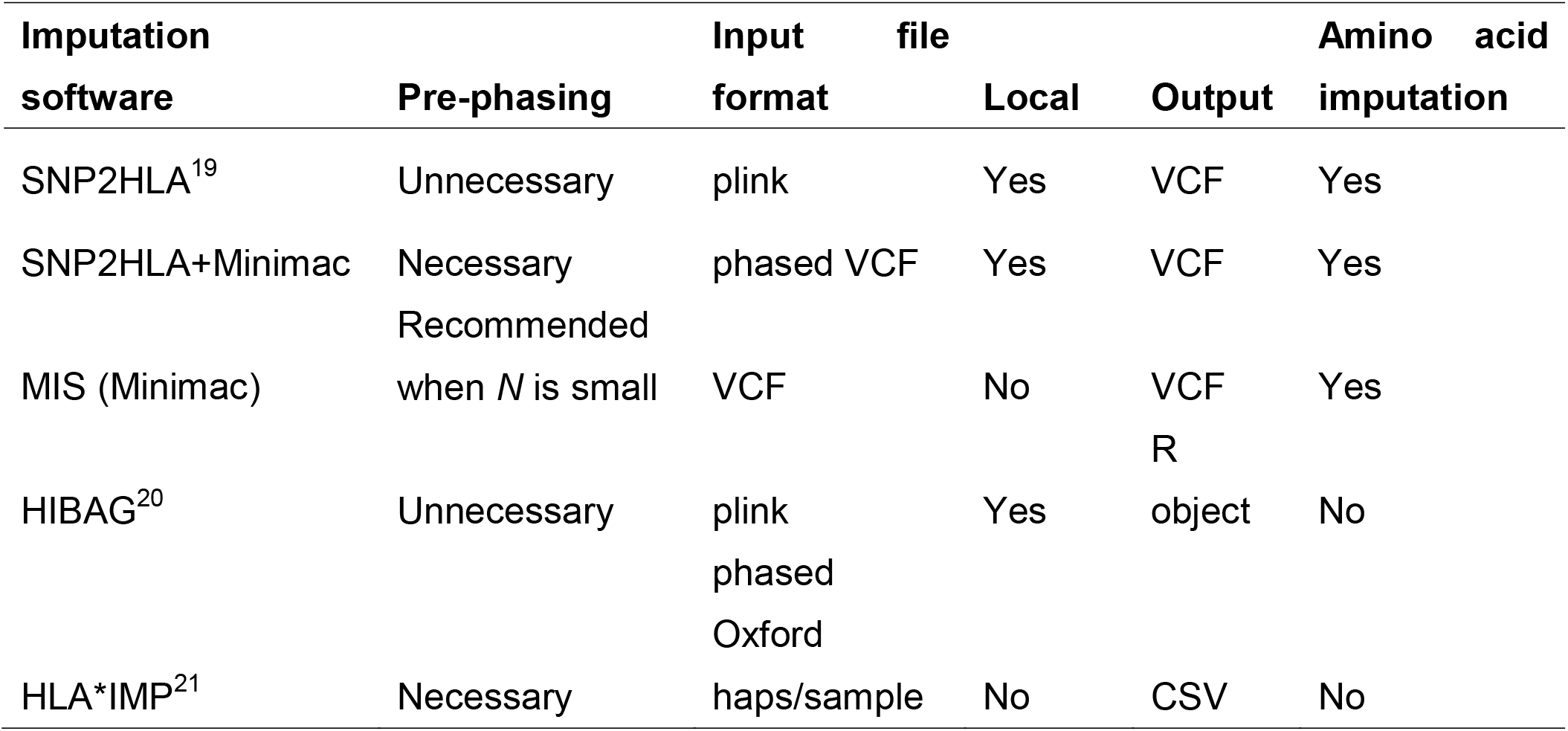
Representative software programs for HLA imputation and their requirements. A list of HLA imputation software programs and their specifications and details about the input and output. MIS: Michigan Imputation Server.

Since our group has developed one of the most widely-used algorithms, SNP2HLA, and its extensions^10,48^, we will focus on the HLA imputation by using the SNP2HLA algorithm along with cloud based implementation at the MIS (URL: https://imputationserver.sph.umich.edu/index.html).

#### SNP2HLA

The SNP2HLA^19^ program can phase and impute HLA alleles, amino acids and intragenic SNPs with HMM implemented in BEAGLE^49^ by taking the QCed target genotype file in the PLINK format as an input. The input file is internally processed to extract variants within the MHC (29 Mb to 34 Mb), and then to correct or remove strand errors when possible based on genotype and AF of palindromic SNPs. In addition to the original bash scripts (URL: http://software.broadinstitute.org/mpg/snp2hla/), there are several extensions to the SNP2HLA algorithm such as HLA-TAPAS^10^ and CookHLA^48^. We also provide a step-by-step explanation of the SNP2HLA implementation, along with a script that allows users to specify all the QC thresholds as option parameters to handle various target cohorts (e.g., the target populations, the number of samples, etc.) in our tutorial website (https://github.com/immunogenomics/HLA_analyses_tutorial).

We note that the original implementation using BEAGLE does not scale to a large number of samples in the target dataset, especially *N* > 10,000. To address this, we also provide a pipeline using the other representative imputation software, Minimac^11^, which can scale to hundreds of thousands to millions of individuals (https://github.com/immunogenomics/HLA_analyses_tutorial). To use Minimac for imputation, we first pre-phase the genotype by using methods such as SHAPEIT^50^ or EAGLE^51^. EAGLE has an advantage of accurate and fast phasing when the number of samples is large (e.g., *N* > 10,000). The pre-phased output file must be converted into the VCF format, and then used as an input to the Minimac software.

#### Michigan Imputation Server

While HLA imputation using the SNP2HLA algorithm can be conducted locally using publicly available HLA reference panels, not all the HLA reference panels are available due to data sharing and privacy restrictions. Our latest multi-ancestry HLA reference panel is one such restricted-access panel^10^. To enable widespread access, we implemented HLA imputation on the Michigan Imputation Server (MIS; https://imputationserver.sph.umich.edu/index.html), which is a cloud-based imputation server with a user-friendly interface (**Supplementary Figure 2**). We host the multi-ancestry HLA reference panel at the MIS and implement the HLA imputation using Minimac as described above. In brief, the user first creates an account online, and securely uploads either a phased or unphased VCF-format genotype file. If the uploaded genotypes are unphased, the uploaded genotype file will be phased within the MIS using the EAGLE algorithm. As noted above, we recommend to pre-phase the genotype (with the reference haplotype when possible) using SHAPEIT or other software when the sample size is small (e.g., *N* < 5,000) to achieve accurate phasing before imputation. The MIS automatically performs basic QC of the input VCF file for the strand orientation and alleles in accordance with the reference. If the input passes the QC steps, the MIS seamlessly performs the HLA imputation. The user will be notified with a download link for the imputed VCF file encrypted with a one-time password via an email once the imputation is completed. The MIS has been used to impute more than 6 million genomes since we started the web-based HLA imputation service in 2021. We benchmarked the performance of HLA imputation on the MIS using individuals with both SNPs and (masked) gold-standard HLA alleles from the 1000 Genomes Project. We confirmed that the imputation accuracy measured by dosage correlation with true HLA alleles was very high across populations (mean dosage correlation *r* = 0.981 for two-field alleles with MAF > 0.05; **Figure 4**).

**Figure 4.**
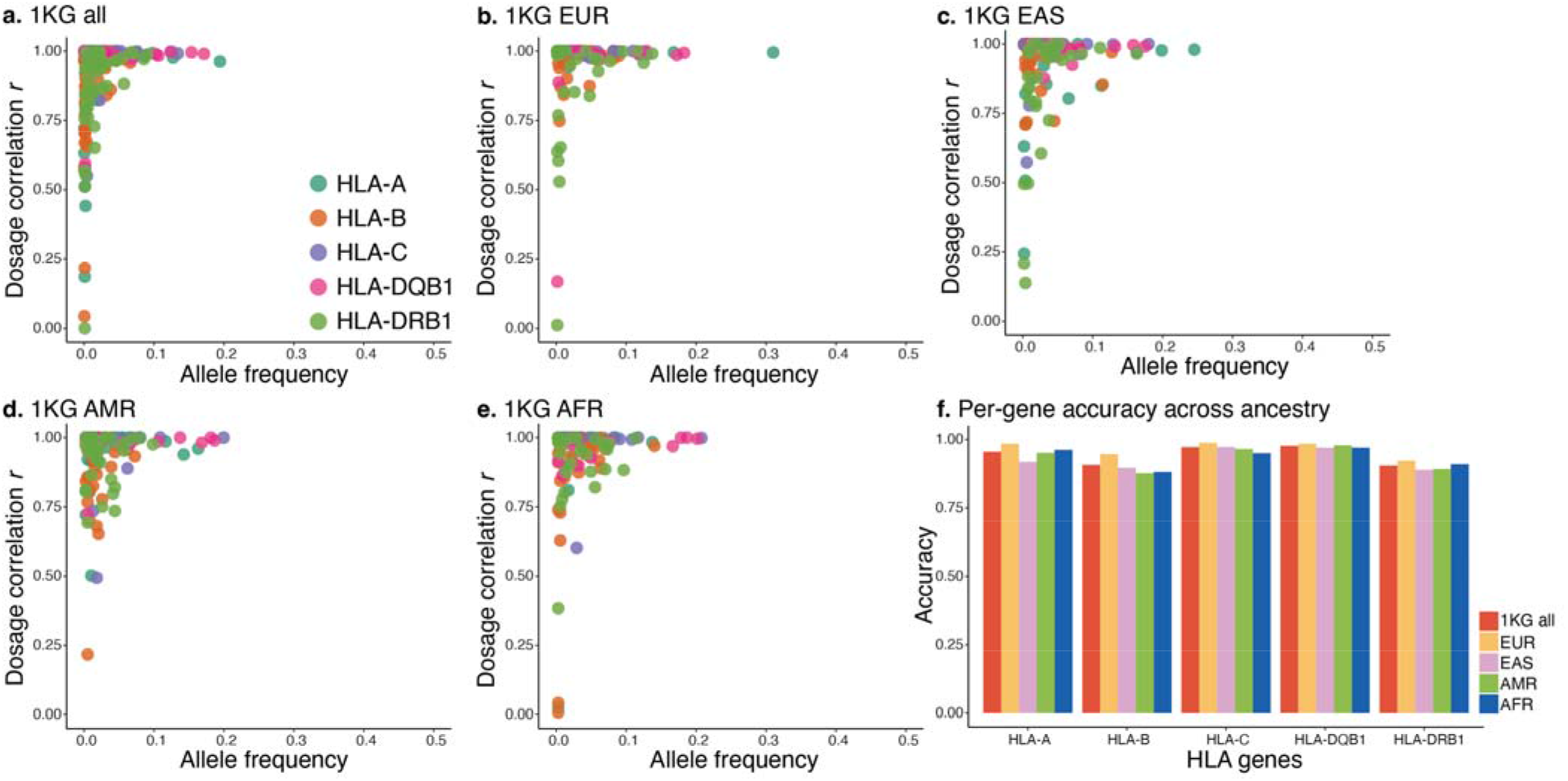
HLA Imputation quality in Michigan Imputation Server. **a**-**e**. Dosage correlation *r* (y-axis) between the Michigan Imputation Server imputed dosage and true genotypes of all two-field alleles in 1KG samples as a function of allele frequency (x-axis), colored by *HLA* gene, for all 1KG individuals (**a**) as well as per-ancestry (**b**-**e**). **f**. The accuracy (concordance) of the imputed dosage of all two-field alleles in1KG samples in Michigan Imputation Server and the true genotype of those per *HLA* gene and per ancestry. The accuracy metric was calculated as previously described^19^.

### Post-imputation QC

The output from the HLA imputation software is accompanied by a quality metric conveying the confidence or estimated accuracy of imputation per allele. A thorough review of these imputation metrics and their correspondence to imputation accuracy is described in Marchini and Howie^15^. We typically QC the imputed HLA alleles, amino acids, and intragenic SNPs based on imputation metrics before association testing. SNP2HLA, Minimac, and MIS all include *Rsq* as a quality metric. The appropriate *Rsq* threshold for QC may depend on the study design; for example, we commonly use *Rsq* > 0.7 in single cohort studies and *Rsq* > 0.5 in multi-cohort meta-analyses. By removing imputed alleles that are below this *Rsq* threshold, some individuals might end up having an *HLA* gene for which the total number of two-field alleles does not sum up to exactly 2. Those individuals might bias the fine-mapping of disease-causing alleles, which we will explain in the subsequent sections. Thus, we recommend removing any individuals that do not have two two-field alleles for a given gene when conducting conditional haplotype tests using two-field alleles.

We recommend calculating true imputation accuracy from classical HLA typing if it is available for a subset of study individuals. While the estimated imputation accuracy generally corelates well with the true accuracy, having the ability to internally benchmark with classical typing for a subset of the cohort is useful for evaluating the true imputation performance, especially if the reference panel imperfectly represents the genetic ancestry of the imputed cohort.

### HLA association and fine-mapping

#### Single-marker tests

Single-marker genetic association tests are used to investigate whether a specific HLA allele, amino acid or SNP is statistically associated with a risk of a disease or a trait of interest, such as risk for a given disease. Similar to the approach used in GWAS, we perform a logistic regression (for case-control traits) or a linear regression (for quantitative traits) for the imputed binary makers that indicate the presence (coded as T in the imputed VCF file) or absence (coded as A in the imputed VCF file) of the selected HLA allele, an amino acid, or an intragenic SNP. For the markers, we typically use the imputed probabilistic dosage genotypes to account for any imputation uncertainty. We include study-specific covariates that could independently explain the trait of interest, such as sex, age, and genotype batches, as well as genotype principal components (PCs) to account for population stratification and an indicator variable of cohorts when combining multiple cohorts.

The logistic regression can be formulated as:

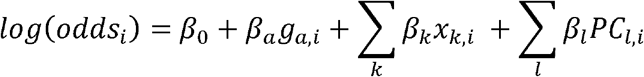

where *log*(*odds*_*i*_) is the logged odds ratio for case-control status in individual *i, a* indicates the specific allele being tested, and *g*_*a,i*,_ is the imputed dosage of allele *a* in individual *i*. The allele *a* could be either HLA alleles, amino acid polymorphisms or SNPs. The *β*_*a*_ parameter represents the additive effect per allele. For all covariates *k, x*_*k,i*_ and *β*_*k*_ are the covariate *k*’s value in individual *i* and the effect size for the covariate *k*, respectively. Similarly,*PC*_*i,i*_, and *β*_*l*_ are the first th genotype PC value in individual *i*and the effect size for the first *l*th genotype PC, respectively, to control for genetic ancestry. The *β*_0_ is the logistic regression intercept.

Quantitative traits that follow continuous distributions (e.g., antibody levels, blood cell counts etc.) can be analyzed by using linear regression similarly:

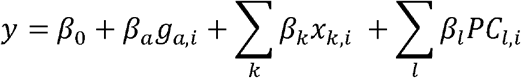

where *y* is a quantitative trait of interest, and normalized by Z score or inverse-normal transformation when necessary.

These association tests can be conducted using conventional GWAS software, such as PLINK^52^, SAIGE^53^, BOLT^54^, etc. by directly using the output VCF files from either the SNP2HLA or the MIS. We use the dosage values designated as “DS” in the imputed VCF files to conduct dosage-based association tests. We provide example command-line scripts to perform single marker tests by using PLINK2 software at our website.

To interpret the results from such an association analysis, we ensure that the “effect allele” (i.e., the allele to which the effect estimate refers) is the presence (coded as P or T in SNP2HLA) of the allele. Also, we note that the association of rare alleles might be spurious due to both the limited accuracy in imputation and the noise in the estimate in the regression. Thus, we might QC the association statistics by MAF to exclude rare alleles (e.g., MAF < 1%). The odds ratio (OR) calculated from the beta (*e*^*β*^) is the estimated risk explained by having one copy of the HLA allele of interest, and the *P* value indicates its significance. Given the strength of LD in the MHC region, trait associations to multiple HLA alleles, amino acid polymorphisms or intragenic SNPs may yield significant results. Further analysis is then required to identify which allele(s) most significantly explains the disease risk within the HLA region.

#### Omnibus tests for fine-mapping amino acid position

To further narrow down the causal position within amino acid sequences within that *HLA* gene, we perform an omnibus test. This analysis is particularly useful when we seek to define mechanisms for the HLA association with the disease, for example by changing the amino-acid compositions at the peptide binding groove of the HLA molecule. In the omnibus test, we estimate the total effect on our trait of interest of all amino acid content variation at a given amino acid position, rather than the separate effects of individual amino acids that appear at that position, as we did in the single-marker test. For an amino acid position which has *M* possible amino acid residues, we assess the significance of the improvement in fit for the full model which includes *M* − 1 amino acid dosages as explanatory variables when compared to a reduced model without including those amino acid dosages. We usually select one amino acid residue that is most common in the studied cohort as the reference allele, and use all the other amino acid residues (*M* − 1) as the explanatory variables. We assess the improvement in model fit by the delta deviance (sum of squares) using an F-test with *M* − 1 degrees of freedom and derive the statistical significance of the improvement.

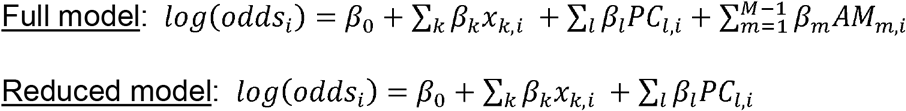

where *m* is one amino acid residue at this position, *M* is the total number of observed amino acid residues at this position, *AM*_*m,i*_ and *β*_*m*_ are the amino acid dosage of the residue *m* in individual *i* and the effect size for the residue *m*, respectively.

We may use the permutation procedure to determine whether the observed association at a single-marker test is primarily driven by HLA alleles (e.g, HLA-DRB1*04:01) or amino acid polymorphisms (e.g., HLA-DRβ1 positions 11, 71 and 74)^12^. To do so, we shuffle the correspondence between amino acid sequences and each of the two-field HLA alleles which was originally defined in IMGT database as described above, while preserving the relationship between the phenotype and the two-field HLA alleles. Then, in each permutation, we select each amino acid polymorphism and assess the improvement in deviance after including this amino acid polymorphism into the model. We typically perform > 10,000 permutations. If the observed improvement using the actual data is significantly larger than the improvements using these permutations, we can infer that amino acid polymorphism is driving the signal, instead of observing the “synthetic” association driven by the HLA allele and its linkage with the causal amino acid(s).

#### Conditional haplotype tests to define a risk sequence of amino acids

Defining the exact stretches of HLA amino acid sequences driving the association with disease allows us to understand the mechanism by which amino acid change affects disease risk ^12^. Importantly, to model combinations of positions, we must use phased genotyping information, rather than encoding each position separately. We perform a conditional haplotype test, where we utilize and combine the imputation results of both two-field alleles and amino acid polymorphisms to obtain phased information. Specifically, we start from the most significant position of amino acid sequence based on the omnibus test we described in the previous section. If there are *M* possible amino acid residues at this position, we can group all possible two-field alleles for this *HLA* gene into *M* groups based on the amino acid residue property at our selected position **(Figure 5a)**. Recall that each two-field allele at a given *HLA* gene corresponds to a unique sequence of amino acids in this gene. In the same way as we did in the omnibus test based on the *M* <u>amino acid residues</u>, we can estimate the effect of each of the *M* <u>groups</u> using a logistic regression model (including covariates, as described above) and derive the improvement in model fit over a reduced model without including those *M* groups.

**Figure 5.**
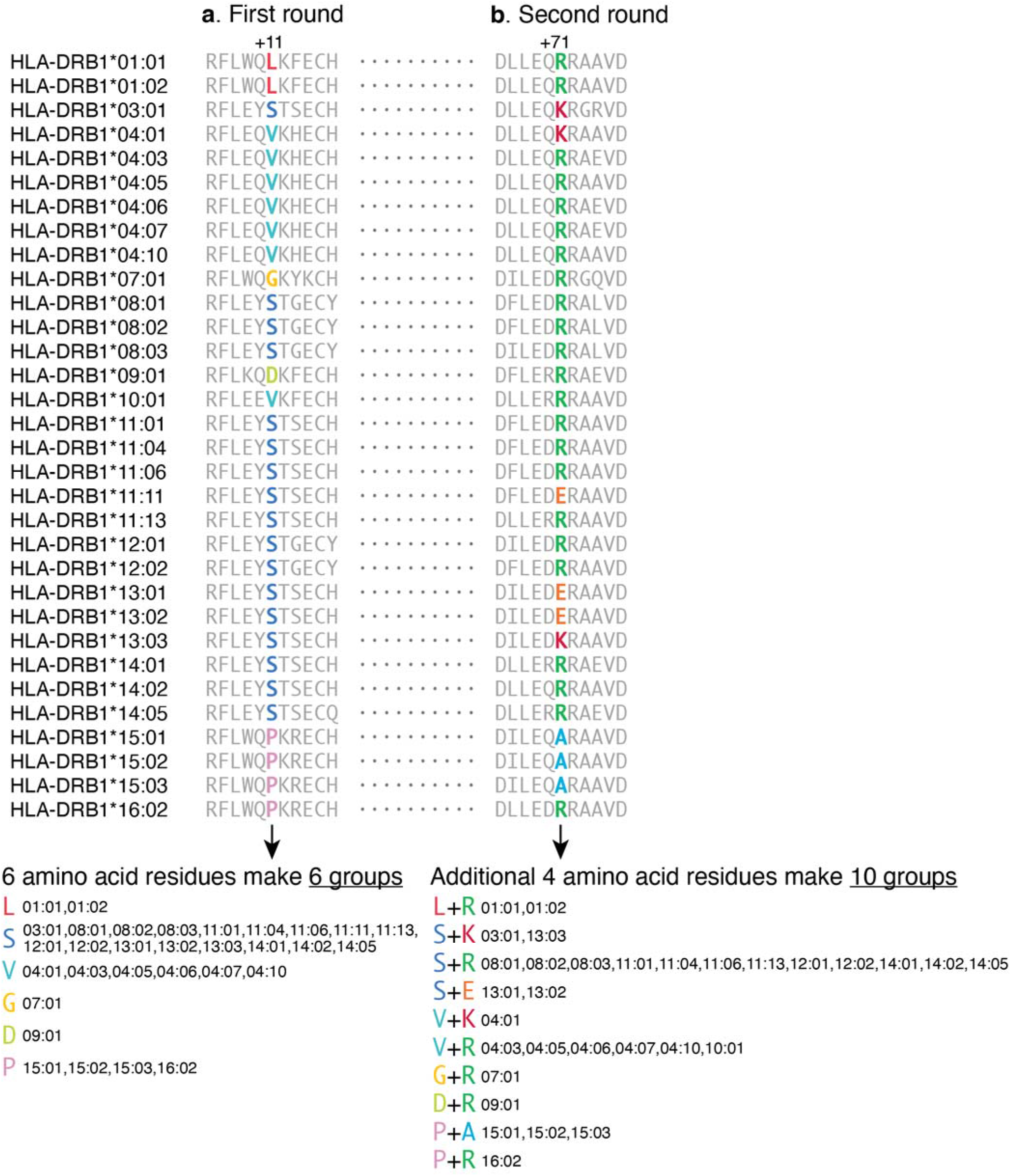
Grouping of two-field alleles in conditional haplotype test. An example illustration of the conditional haplotype test for the *HLA-DRB1* gene. In the first round of the amino acid association test at position +11 (**a**), we group all two-field alleles (32 alleles in total) into 6 groups based on the amino acid residues at the position +11, and ask whether those groups significantly explain the disease risk by using omnibus test. In the second round of conditional haplotype test (**b**; position +71 as an example), we group the two-field alleles into 10 groups based on the amino acid residues at the position +11 and +71. Then, we ask whether those 10 groups explain the disease risk more significantly than the 6 groups that we defined in the first round.

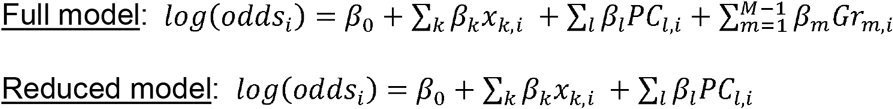

where *Gr*_*m,i*_ is the sum of the dosage of two-field alleles from a group *m*, explained by the *m*’th amino acid residue. We note that we recommend removing any individuals that do not have two two-field alleles for a given gene, as we explained in the *Post-imputation QC* section. Once we define the most significant individual position at a given *HLA* gene based on the significance of improvement, we next seek to identify which amino acid position other than this significant position best improves the model over the model only including this significant position (**Figure 5b**). Let *x* be the most significant position in the primary analysis, which has *X* possible amino acid residues. We sequentially test each amino acid position (*z*) other than *x*, to ask whether haplotypes defined by the amino acid combination of positions *x* and *z*(*z* ≠ *x*) explain the disease risk more than those defined only by the position *x*. To do so, we re-categorize all two-field alleles at this *HLA* gene into *Z* groups, where *Z* is the total number of observed haplotypes defined by the amino acid positions *x* and *z*. The value of *Z* must be at least *X* if no new haplotypes are defined. We again assess the significance of the improvement in model fit of the Full model (covariation at positions *x* and *z*) over the Reduced model (variation at position *x* alone) by the delta deviance (sum of squares) using an F-test with *Z*−*X* degrees of freedom.

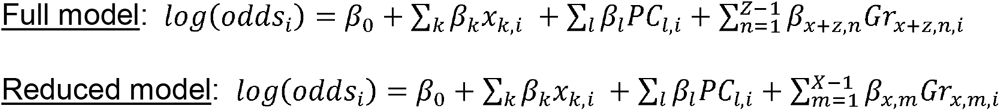

where *Gr*_*x*+*z,n,i*_ is the sum of the dosages of two-field alleles in a group *n* by a given combination of the amino acid residues at positions *x* and *z*.

Thus, we define the next most significant amino acid position which additionally and independently explains the disease risk from the position *x*. If the model improvement in this second round is statistically significant, we iterate the same analyses to identify amino acid position(s) other than the previously identified positions that best improve the model over the model including those previous positions, until we obtain no further significant improvement from any of the remaining positions.

#### Tests for non-additivity

The dosage effect of HLA (having one copy or two copies of a given HLA allele) on disease risk is not purely additive in infectious diseases and autoimmune diseases^55–63^. All the analyses we have described above assume the additive risk model, a model in which the risk (i.e., log Odds Ratio(OR)) for acquiring a disease due to carrying one copy of the allele (heterozygous state) is half the risk (log(OR)) conferred by carrying two copies (homozygous state). A non-additive effect represents a deviation from this linear relationship between the dosage and the risk (**Figure 6a**). For instance, a dominant effect might be indicated when the effect of carrying one copy is more than half the effect of carrying two copies. A biological explanation for such a dominant effect might be (1) having one copy is enough to express the MHC variant with the disease-relevant antigen-binding properties on the cell surface, or that (2) there are synergistic interactions with another HLA allele at the same locus. Lenz et al.^62,64^ showed that such non-additive effects are pervasive in a spectrum of autoimmune diseases.

**Figure 6.**
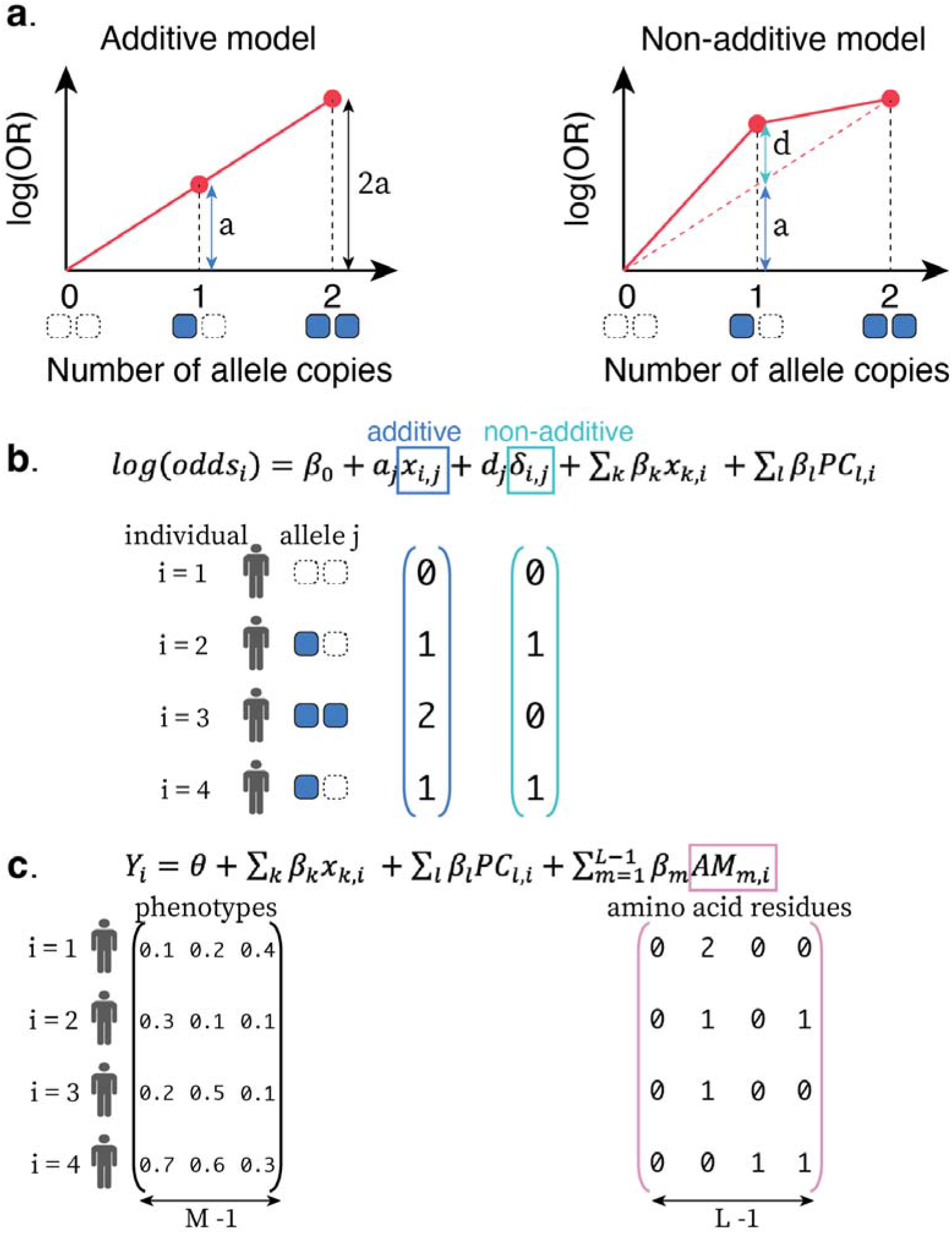
Non-additive test and multi-trait analysis. **a**. Schematic illustrations of additive model and non-additive models using the log odds ratio (log(OR)) according to the dosage of the genotype of interest. *a* denotes the purely additive effect by having one copy of the allele, and *d* denotes any departure from additivity at heterozygous genotype. **b**. A logistic regression model to assess both the additive and non-additive effect of the allele *j* (see main text for details). **c**. Multi-trait analysis by using multiple linear regression model (MMLM) to test the association between multi-dimensional phenotype *y* and the amino acid polymorphism.

To test for the non-additive effect, we construct a logistic regression model which captures both additive and non-additive contribution of the allele to the disease risk (**Figure 6b**)^62,65^. We first define the additive term *x*_*i,j*_ as either the best guess or the dosage genotype of allele *j* in an individual *i* which we are interested in.

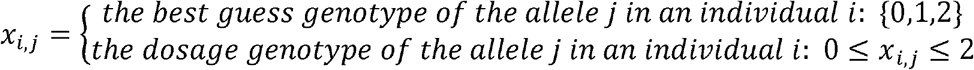

We next define the non-additive term *δ*_*I,j*_ as the heterozygous status of the allele *j* in an individual *i*, which should capture any deviation of the effect from the additivity.

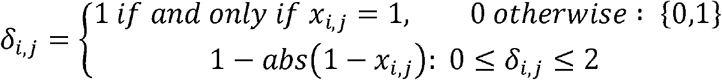

Using those two terms *x*_*i,j*_ and *δ*_*i,j*_, we construct a full model by including both additive and non-additive term with covariates, and a reduced model with by including only additive term with covariates.

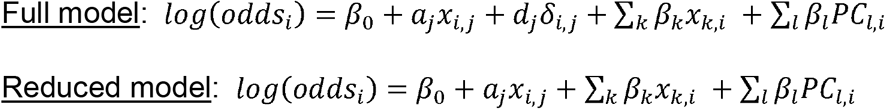

where *a*_*j*_ denotes an additive effect and *d*_*j*_ denotes a non-additive (dominance if positive) effect.

We finally assess the significance of the improvement in model fit of the Full model over the Reduced model in model fit by the delta deviance (sum of squares) using an F-test.

#### Tests for interactions among HLA alleles

Once we identify an allele harboring a possible non-additive effect, we may also be interested in understanding whether this is due to an interaction effect between the identified allele and the other allele at the same HLA locus. In other situations, we may want to assess an interaction effect between a pair of alleles of functional interest. If the disease risk from a combination of those two alleles deviates from the expected disease risk by multiplying the disease risk (i.e., adding the log(OR)) of each of the two alleles, that combination can be regarded as having an interaction effect. To test this hypothesis, we construct a reduced model which only includes an additive term for each of the two alleles, and a full model which includes an interaction term between the two alleles in addition to the additive term for each of the two alleles. Let *x*_*i,j*_ be the dosage genotype of the allele *j* in a given individual *i* nominated by a significant non-additive test, and let *x*_*i,h*_ be the dosage genotype of the other allele *h* (*h* ≠ *j*) in an individual *i* to be tested for an interaction effect with the allele *j*.

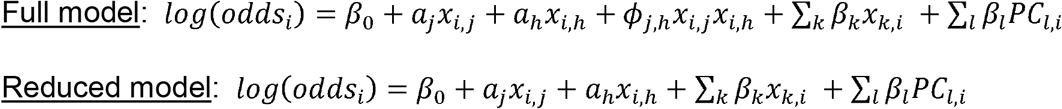

where *ϕ*_*j,h*_ is the effect size of the interaction between the alleles *j* and *h*. We again assess the significance of the improvement in Full model over Reduced model in model fit by the delta deviance (sum of squares) using an F-test. We note that the observed interaction effects can be spurious when the frequencies of the tested alleles are relatively low, which results in noisy effect estimate. We consider conservative QC of the tested alleles based on MAF (e.g., MAF > 0.05 or 0.10), or performing permutation analyses to test whether the observed statistics could occur by chance, in such cases.

#### HLA evolutionary allele divergence

A potential source for non-additive interaction effects among HLA alleles is the extent to which their encoded HLA molecule variants differ functionally (i.e., in their bound antigen repertoires). Since HLA genes are generally co-dominantly expressed, both HLA variants of a heterozygous individual are presenting antigens at the cell surface. If two HLA alleles are very similar in their sequence, their encoded HLA molecules on average will bind similar sets of antigens and thus exhibit a substantial overlap in their presented antigen repertoires, while the opposite will be true for two alleles with very divergent sequences^66^. The concept that carrying two divergent HLA alleles will allow HLA-presentation of a wider range of antigens, and by extension increase the likelihood of pathogen detection by the adaptive immune system, has been termed *divergent allele advantage* (DAA), as an extension of the classical heterozygote advantage^67,68^. DAA has already been shown to drive HLA allele frequencies and contribute to HIV control^63,66^. but might have broader implications in HLA-mediated complex diseases. For instance, it was shown that cancer patients whose HLA class I alleles exhibit a higher HLA evolutionary divergence (HED) respond better to cancer immunotherapy, possibly because more mutated neoantigens are presented by their HLA^69^. The HED score between two HLA alleles at a given HLA locus is based on the Grantham distance between their amino acid sequences, The HED is applicable to both HLA class I and class II alleles. It can be calculated using publicly available scripts^66^, and its effect on a given phenotype can then be estimated by adding it as a quantitative parameter in a regression model and testing for improvement in model fit with an F-test.

#### Multi-trait analysis

Our group recently showed that the amino acid frequencies at complementarity-determining region 3 (CDR3) of the T cell receptor (TCR) are highly influenced by the HLA alleles and amino acids, possibly through thymic selection^6^. This type of analysis is an extension of the analyses we described in the previous sections. One notable difference is that the response variable represents not a single trait (e.g., a disease) but multiple traits: in this case the frequencies of each amino acid residue at the position of interest within CDR3, which we call cdr3-QTL analysis. We test which amino acid position has a significant association with those frequencies overall, using an extended framework of the omnibus test that we described above (**Figure 6c**). In this case, the response variable is not a vector of one phenotype, but a matrix (multidimensional vector) of frequency phenotypes where each row represents an individual and each column represents a frequency of a given amino acid residue at a given position of CDR3. Let *Y* be this frequency matrix with *N* rows and *M* − 1 columns, and *Y*_*i*_ be the *M* − 1 frequency phenotypes in an individual *i*,. *N* denotes the number of individuals, and *M* denotes the number of observed amino acid residues at this position. We perform a multivariate multiple linear regression model (MMLM) to test the association between *Y* and HLA alleles or amino acid positions of interest.

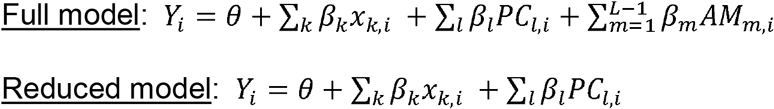

where *θ* is an M-dimensional parameter that represents the intercept, *L* is the total number of observed amino acid polymorphisms at this position, *AM*_*m,i*_, and *β*_*m*_ are the amino acid dosage of the residue *m* in an individual *i* and the *M*-dimensional effect sizes for the residue *m* on *Y*, respectively.

We assess the significance of the improvement in model fit between Full model and Reduced model with the multivariate analysis of variance (MANOVA) test for quantitative traits. As spurious associations again arise when the frequencies of the tested alleles are relatively low ^6^, we recommend performing permutation analyses to confirm the calibration of the test statistics.

By using this multi-trait framework, we can assess any combination of multiple phenotypes. One potential application is to investigate disease phenotypes by using deep phenotype record in biobanks. This framework could disentangle pleiotropic HLA alleles that simultaneously affect a spectrum of diseases of interest. Another interesting application might be multiple molecular phenotypes such as expression or protein abundance of multiple genes, and a combination of multiple modalities (e.g., expression and chromatin accessibility). We can also assess those phenotypes across multiple cell types (e.g., expression of a gene in T cells, B cells, Monocytes etc.).

#### Concluding remarks

Given the increasing number of associations between the HLA region and human complex traits that have been identified through large-scale GWAS, accurate imputation and fine-mapping of the causal HLA alleles and amino acids will continue to be important as the data size continues to grow. We present a strategy that can lead investigators to fine-mapped alleles. Leveraging HLA fine-mapped alleles with the variants outside of MHC region, it may be possible to construct an efficient genetic risk score to stratify people based on the genetic risk for those diseases. We have publicized this imputation pipeline through the user-friendly MIS, which hosts the HLA reference panel representing multiple populations and enables web-based automatic HLA imputation for global cohorts. Another advantage of this implementation is the computational efficiency: HLA imputation of a cohort of millions of individuals is computationally scalable (for example, for a cohort of size 20,000, HLA imputation runs within 1 hour). We hope this protocol will empower the field of statistical genetics to more comprehensively define the effect of HLA variation on a spectrum of human diseases.

Despite the well-established performance of our approach, we can still improve our HLA imputation reference panel further. First, we continue to expand the reference panel to better represent global populations that are currently missing (e.g., Africans and South Asians). Similarly, the scope of genes included in the panel can be expanded to include, for example, non-classical *HLA* genes and *C4* copy number. Second, the imputation accuracy is currently satisfactory in association testing but not yet as high as the gold-standard HLA typing. We aim to further improve the accuracy by updating the HLA calls and scaffold variants used in the reference panel as well as improving the imputation algorithms.

While fine-mapping of HLA alleles has provided deeper insights into disease pathogenesis, we need more mechanistic or structural understanding of how these alleles contribute to disease biology. Why do certain HLA alleles cause a diverse spectrum of diseases? How do those alleles characterize our inherited composition of T cell repertoires? What are auto-antigens that are being presented by those alleles? Recent advances in experimental and computational modeling of protein structures and its complex^70,71^ can offer promise. We need both experimental and computational approaches to answer all these important questions.

## Data Availability

We provided the availability of HLA imputation reference panel at Table 1. We made our HLA imputation pipeline using multi-ancestry HLA reference panel publicly available at Michigan Imputation Server (https://imputationserver.sph.umich.edu/index.html).

## Code Availability

The computational scripts and their usage related to this tutorial are available at https://github.com/immunogenomics/HLA_analyses_tutorial.

## Acknowledgments

This work is supported in part by funding from the National Institutes of Health (R01AR063759, U01HG012009, UC2AR081023). S.Sakaue was in part supported by the Manabe Scholarship Grant for Allergic and Rheumatic Diseases and the Uehara Memorial Foundation. J.B.K. was supported by NIH/NIGMS T32GM007753. A.J.D. was funded by NIH/NIDDK T32DK007028. T.L.L. was funded by the Deutsche Forschungsgemeinschaft (DFG, German Research Foundation) – Projektnummer 437857095.

## Author Contributions

S.Sakaue and S.R. conceived the work and wrote the manuscript with critical input from all authors. S.Sakaue and S.G. created a web tutorial accompanying this manuscript. All authors contributed to developing this protocol. S.Sakaue, M.C., Y.L., W.C., S.Schönherr, L.F., J.L., C.F., A.V.S., and S.R. contributed to updating the multi-ancestry HLA reference panel and implementing HLA imputation at Michigan Imputation Server.

## Competing Financial Interests

B.H. is a CTO of Genealogy Inc. T.L.L. is a co-inventor on a patent application for using HED in predicting cancer immunotherapy success.

## Notes

https://github.com/immunogenomics/HLA_analyses_tutorial

