## Supplementary Figures and Tables for "A statistical genetics guide to identifying HLA alleles driving complex disease"

**Sakaue et al.**

**Table of contents:**

Page 3	**Supplementary Figure 1**

Page 4	**Supplementary Figure 2**

Page 5	**Supplementary Table 1**

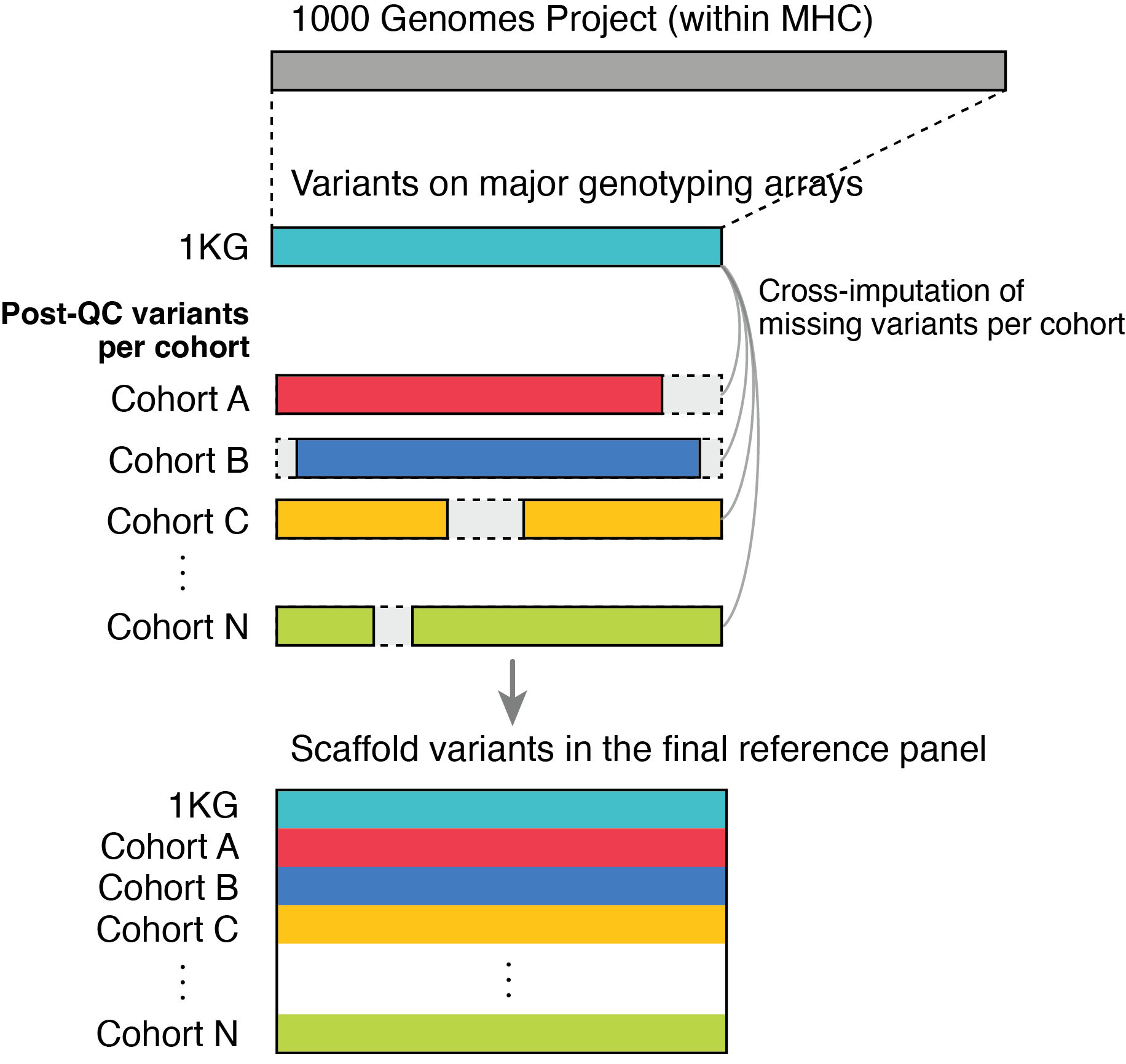

**Supplementary Figure 1 | Schematic illustration of method used to construct scaffold variants within multi-ancestry HLA reference panel.**

We extracted SNP variants within MHC region in 1000 Genomes Project (1KG) samples. We only retained variants that were included in major genotyping arrays (Illumina Multi-Ethnic Genotyping Array, Global Screening Array, OmniExpressExome, and Human Core Exome), colored in teal. We then QCed each of the participating cohorts’ MHC SNPs separately, retained overlapping variants with selected SNPs in 1KG, and cross-imputed each cohort’s missing variants by using 1KG genotypes. We finally combine all cohorts together to construct multi-ancestry scaffold variants.

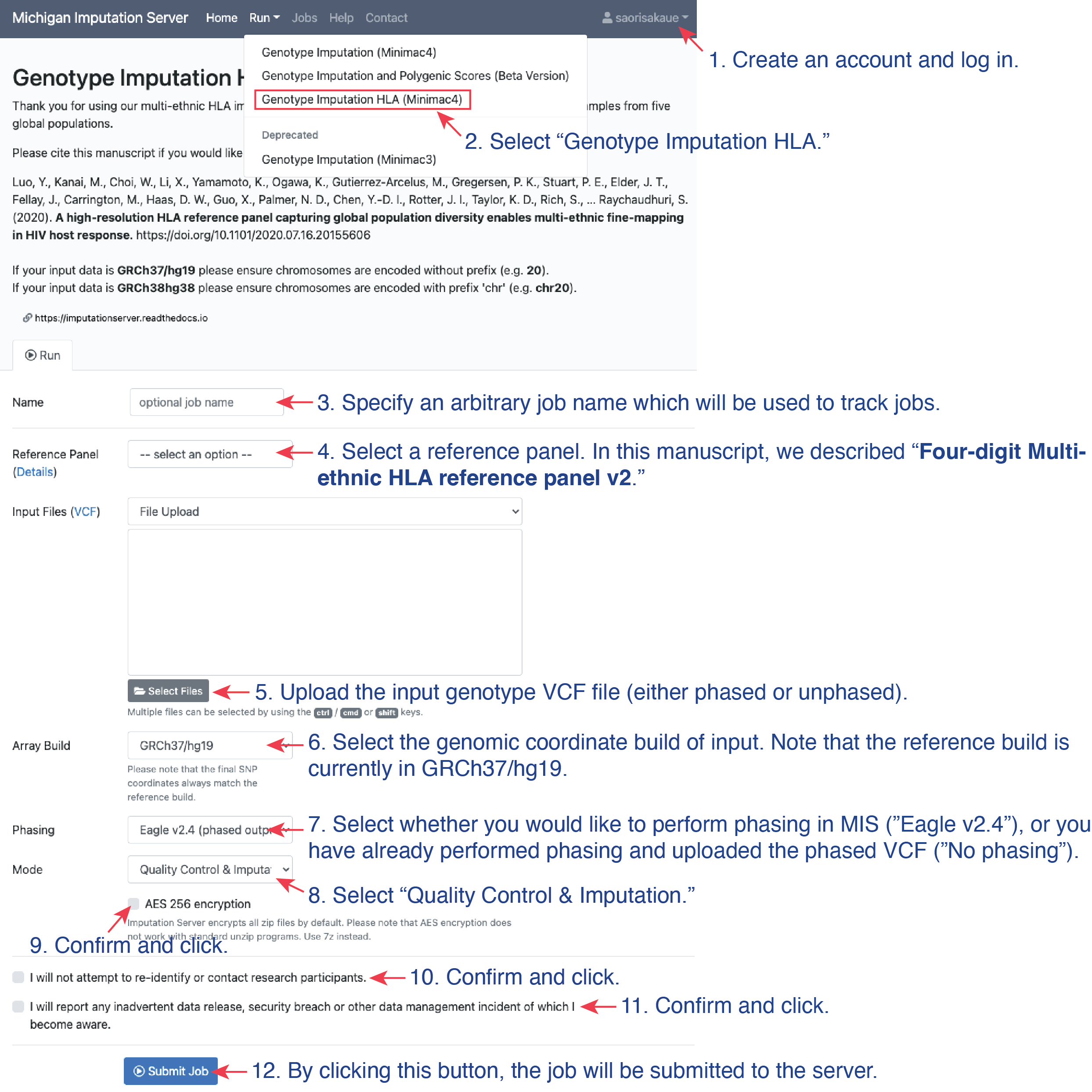

**Supplementary Figure 2 | Michigan Imputation Server.**

Example usage of Michigan Imputation Server for HLA imputation at https://imputationserver.sph.umich.edu/index.html

| **Array** | **N_in_MHC_** | **N_in_HLA_panel_** |
| --- | --- | --- |
| Affy6.0 | 2,433 | 1,805 |
| Axiom_AveraNTR.na35 | 8,958 | 6,708 |
| Axiom_GW_ASI_SNP.na34 | 7,546 | 4,770 |
| Axiom_GW_CHB2.na34 | 2,906 | 1,526 |
| Axiom_GW_EUR.na34 | 9,511 | 5,484 |
| Axiom_GW_LAT.na34 | 9,817 | 5,855 |
| Axiom_GW_PanAFR.na34 | 3,818 | 1,837 |
| Axiom_PMRA.na35 | 9,197 | 7,003 |
| Axiom_UKB_WCSG.na34 | 10,436 | 7,190 |
| cytosnp-850k_b | 6,695 | 5,316 |
| DrugDevConsortium_15073507_A1 | 3,257 | 1,448 |
| GSA-24v3-0_A1 | 8,017 | 7,437 |
| GSAMD-24v1-0_20011747_A4 | 8,783 | 7,703 |
| human660w-quad_v1_h | 2,604 | 1,951 |
| humancore-12v1-0_a | 1,080 | 858 |
| HumanCytoSNP-12v2-1_H | 712 | 632 |
| humanomni2.5-4v1_h | 9,742 | 6,373 |
| humanomni5-4v1_c | 33,371 | 11,787 |
| humanomniexpress-12v1-1_b | 6,434 | 5,294 |
| HumanOmniZhongHua-8-v1-0-C | 10,918 | 7,490 |
| InfiniumExome-24v1-1_A1 | 3,905 | 2,370 |
| InfiniumImmunoArray-24v2-0_A | 9,233 | 8,698 |
| Multi-EthnicAMR-AFR-8v1-0_A1 | 14,138 | 10,539 |
| Multi-EthnicEUR-EAS-SAS-8v1-0_A1 | 14,507 | 10,577 |
| Multi-EthnicGlobal_A1 | 15,762 | 11,730 |
| OncoArray-500K_B | 5,974 | 3,610 |
| PMDA.hg19 | 12,233 | 8,493 |
| PsychArray-B | 4,779 | 3,097 |

**Supplementary Table 1 | The number of variants within MHC (28–34Mb) and the number of SNPs overlapping with our HLA imputation panel.**

The selection of the arrays and variant information is based on <https://doi.org/10.1038/s41431-021-00917-7>. Raw data was downloaded from https://github.com/jverlouw/ArrayComparisonData and we assessed those numbers based on the rsID information.
